# Predictive perception via simultaneous learning and inference

**DOI:** 10.64898/2026.09.24.754081

**Authors:** Mahdi Enan, Mario Senden, Paulina Janik, Yamil Vidal, Ryszard Auksztulewicz, Federico De Martino

## Abstract

Perception has been proposed to involve an inferential process that combines noisy sensory evidence with prior expectations to estimate the latent states of the environment. Exact Bayesian inference is intractable in the continuous, high-dimensional state spaces of the natural world. Variational schemes address this by restricting the form of the posterior, whereas sampling schemes represent the posterior with a finite set of samples. In both cases the approximation concerns the posterior rather than the state space over which it is defined. Here we place the approximation elsewhere. Rather than restricting the form of the posterior, we restrict the state space, modeling environmental states as discrete and thereby making Bayesian filtering exact. Beliefs are then unconstrained in shape and what has to be learned can be reduced to a transition matrix updated online by a local rule. The cost is that the states, rather than the distribution, must be specified in advance. The Simultaneous Learning and Inference Model (SLIM) combines this exact filtering with a gated Hebbian rule that learns transitions from inferred rather than observed states. In simulations SLIM recovers environmental dynamics under sensory noise and adapts when those dynamics change, with error growing only once the sensor becomes uninformative. A hierarchical instantiation reproduces local and global prediction error effects. Applied to two auditory decision-making experiments under noise, SLIM reproduces behavioral signatures of expectation and supplies trial-level measures of expectation and surprise. Exact inference over a small discrete state space with a single local Hebbian rule is therefore sufficient to account for these phenomena, without an explicit optimization objective and without a parametric approximation to the posterior.

## Introduction

Humans can effortlessly process sensory stimuli in noisy environments, leveraging computational principles such as probabilistic inference, yet the implementations underlying this capability are not well established [1]. The Bayesian brain hypothesis proposes that the brain achieves stability in the face of noise and uncertainty through probabilistic inference [2]. According to this hypothesis, the brain infers hidden causes of its sensory input by combining prior expectations with noisy observations akin to the application of Bayes’ theorem. Within this perspective, hidden causes are formalized as states of the environment, sensory input governs the likelihood function, and internal expectations encode a prior distribution over states. These are combined to form a posterior distribution (i.e., a belief) once sensory evidence is taken into account [2]. For example, the brain may use sensory information to infer that the environment contains a beat within a piece of music, a word in a sentence, someone’s facial expression during a conversation, or any other phenomenon which can be characterized as a distinct perceivable state within the sequence of states that describe the environment.

While this Bayesian formulation is conceptually appealing, exact inference is generally intractable in the continuous, high-dimensional state spaces characteristic of the natural world. The main reason underlying this intractability is that computing the evidence of a given sensory input (i.e. the denominator of Bayes’ formula) requires integrating over all possible states of the environment, which has generally no analytic solutions in continuous and high-dimensional state spaces. This limitation has led to the development of two major strategies for approximating beliefs, each involving different trade-offs [3]. Variational schemes typically retain a rich state space but restrict the form of the posterior, whereas sampling schemes do not assume a particular posterior form but approximate the posterior with a finite set of samples. In both cases, the approximation concerns the posterior rather than the state space over which it is defined.

The free energy principle (FEP), for example, makes inference tractable by employing a variational scheme in which the approximate posterior is assumed to belong to a restricted family of distributions [4–6]. This approach transforms the inference problem into a mathematical optimization problem. Rather than computing the exact posterior, the brain is assumed to represent a simpler and tractable approximate posterior based on a parameterized family of distributions. While early applications utilizing variational inference focused on modeling perception as inference by approximating continuous posteriors using simple Gaussian distributions [7], more recent work integrates perception and action while capturing multi-modal beliefs using discrete Dirichlet distributions [8]. The free energy principle provides a highly elegant and unified framework that combines perception, action and learning using Markov decision processes. However, tractability is obtained by committing to a specific choice of the variational distribution to be used, by ensuring that posterior beliefs remain in the same family of distributions through a parametrization of the posterior beliefs, and by assuming independence between factors of the approximate posterior beliefs. Ultimately, this places the requirement on the experimenter to pick the appropriate tractable variational distribution family for a given task.

An alternative to variational approaches is sampling-based inference. Particle filters ([9]), for example, achieve tractability by sampling from a predictive prior distribution and weighting each sample with the likelihood of observations to approximate the posterior distribution non-parametrically. These techniques overcome the restrictions posed by assuming a parametrized family of distributions and allow for flexible online Bayesian filtering; that is, recursive updating of probability distributions over time in response to noisy observations and control inputs. However, this flexibility comes with an increased computational cost compared with variational inference. While the original particle filters (which approximate Bayesian filters) lacked biological realism because they rely on operations which do not have clear neuronal implementations (such as importance weighting and resampling), more recent work has shown that biologically plausible implementations are possible [10]. Instead, Kalman filters perform exact analytical Bayesian filtering but assume linear Gaussian state spaces, [11].

Each of these strategies therefore comes with its own trade-offs, and broader problem formulations can be realized with either of them. In control theory and reinforcement learning, for example, partially observable Markov decision processes (POMDP) provide a unified framework which integrates Bayesian filtering of hidden states with decision making under uncertainty. When exact belief updating is computationally intractable, belief states can be approximated by finite sets of particles (samples) using particle filtering. The POMDP framework is also applied in active inference models [12], where beliefs are instead updated variationally. In both cases, however, the standard formulation takes the transition and observation models as given, leaving open how they are learned.

In the brain, learning and inference are not always cleanly separated (see e.g. [4] for a description of the distinction between perception and learning). In perceptual decision-making, sensory evidence is often insufficient to uniquely determine the underlying state of the environment, requiring observers to combine noisy observations with prior expectations. Moreover, inference is not performed in isolation from context: ambiguous cues can alter the interpretation of subsequent sensory inputs by modifying expectations about likely environmental states. Where the states themselves are uncertain, the signal available for updating an internal model of environmental dynamics is not the sequence of observations but the observer’s own beliefs about which states produced them. The two approaches of approximation address this in different ways. Within variational approaches the loop is closed by using the approximate posterior itself as the learning signal. In predictive coding, the weights encoding the generative mapping are updated by the product of a prediction error and the inferred activity at the level above, so what is learned is driven by the causes the model infers rather than by the sensory input directly [6, 13]. What is learned in this way is a set of parameters within an assumed structure, and the update inherits whatever has been assumed about the form of the posterior. Many free energy-based models accordingly assume a predefined generative model structure and focus primarily on inference and parameter updating within it. However, evidence from perceptual learning [14, 15], from the tracking of environmental volatility [16] and from experience-dependent plasticity [17–19] indicates that the brain must continually adapt not only its expectations but also its internal representations of environmental regularities as conditions change. Active inference methods can enable agents to learn discrete internal model structures in small state spaces [20], while deep active inference techniques trade biological realism for scalability to learn in high-dimensional continuous environments using gradient-based optimization [21].

Sampling-based approaches face the same problem. Estimating a model of the environment while simultaneously inferring states from it is a familiar problem in robotics, where simultaneous localization and mapping (SLAM) requires a robot to estimate its own position while learning the map on which that estimate depends [22]. More generally, estimating the parameters of a state-space model from within a particle filter is a mature area of statistics, addressed by particle Markov chain Monte Carlo [23], SMC^2^ [24] and kernel smoothing over parameter particles [25]. These methods retain the asymptotic exactness of sampling, but obtained either offline over a batch of observations [23] or by nesting a second filter over parameters [24]. Genuine online variants, such as kernel smoothing over parameter particles [25], avoid that cost but introduce a bias that is difficult to quantify, while augmenting the state with static parameters is subject to path degeneracy [26].

The trade-off each approach accepts is not limited to the level of inference because what is approximated determines what an implementation has to represent. Where the posterior is restricted to a parametric family, beliefs are summarized by a small number of sufficient statistics. Under Gaussian assumptions the updates to those statistics take the form of local Hebbian-like plasticity, as in predictive coding [6]. When the posterior is instead represented by samples, the distribution is carried by variability itself, which has been proposed as an account of stochastic neuronal responses [27]. Implementations in both approaches nonetheless span a spectrum, from local - biologically plausible mechanisms such as Hebbian plasticity-to scalable optimization-based approaches relying on gradient descent. In both approaches, it is the scalable variants that depend on gradient-based optimization or on batch computation. In biological systems, these complementary mechanisms may be reflected in distinct forms of plasticity, including NMDA-dependent synaptic changes in neocortical circuits [28] and faster learning of predictive representations through recurrent interactions in hippocampal regions such as CA3, potentially supporting the formation of successor representations [29].

While variational and sampling schemes diverge in how they treat inference, learning and neural implementation, what they have in common is more consequential than what separates them. In the continuous, high-dimensional settings from which the problem of intractability arises, both cab retain a rich state space and place the approximation on the posterior defined over it. That commitment is what allows either variational and sampling schemes to account for a wide range of perceptual and cognitive phenomena [4, 30]. It is also where their costs arise, whether in the bias due to commitments made about the form beliefs may take, or in the expense of obtaining enough effectively independent samples. We therefore asked what follows from placing the approximation elsewhere. Rather than restricting the form of the posterior, we restrict the state space, modeling environmental states as discrete and thereby leveraging the tractability of exact inference in finite state spaces [3]. We restrict the state space to enable exact inference, leaving the shape of beliefs unconstrained and reduce the internal representation of environmental dynamics to a transition matrix that can be updated by a biologically inspired local rule. The choice between restricting the form of the posterior and approximating it with samples therefore does not arise, since a belief over a small set of discrete states is simply a vector of probability masses, one for each possible state. The cost is that the states, rather than the distribution, must be specified in advance.

We justify this assumption of discrete perceptual states based on evidence that perception often exhibits categorical organization, including the categorical perception of speech sounds [31] and of melodic musical intervals in trained musicians [32] and the categorical representation of continuous sensory input in auditory and prefrontal cortex [33, 34]. Discrete states can moreover be understood as a coarse-grained description of continuous neural dynamics that settle into a finite number of stable basins, as in bistable perception [35, 36] or in the abrupt transitions between hippocampal representations that are observed when the environment is morphed continuously [37, 38]. Importantly, this is an assumption about the model’s internal representation rather than about the environment itself. The sensory system is assumed to map continuous input onto a discrete state space.

Here we introduce the Simultaneous Learning and Inference Model (SLIM) which obtains tractability by utilizing exact Bayesian filtering within a restricted discrete state space. We demonstrate that in this framework the internal representation of environmental dynamics can be updated online via algorithmically plausible, local Hebbian mechanisms that track state transitions. To achieve this, we propose a gated Hebbian rule that learns discrete model structures from inferred environmental states. The gating serves two purposes. It ensures that the updated representation remains a valid set of conditional probabilities without a separate normalization step, and it restricts each update to the transitions relevant in the current context. Estimating transition probabilities online in this way connects SLIM to accounts of human sequence learning in which such probabilities are tracked directly from observed events [39], with the difference that SLIM estimates them from states it infers and is therefore applicable where the events themselves cannot be unambiguously identified from the sensory evidence. While the brain can easily learn simple relationships, it also possesses a remarkable capacity to learn higher-order dynamics of the environment by processing interactions that evolve over larger timescales [40–43]. Accordingly, our learning rule can be parameterized to enable learning of higher order associations (see Methods section on learning).

Through simulations, we show that SLIM recovers environmental dynamics under sensory noise and adapts when those dynamics change. The recovery has a limit, since the error in the learned dynamics grows once the sensor becomes uninformative and beliefs are driven by the model’s own predictions alone. A hierarchical instantiation of SLIM reproduces the distinct prediction error signatures elicited by local and global regularity violations [44, 45]. We then apply the model to two auditory decision-making experiments conducted under constant background noise, the second of which was designed for the present work and adds an ambiguous cue that allows the effect of expectation on detection to be separated into a benefit for expected targets and a cost for unexpected ones. Fitted to individual participants, SLIM reproduces behavioral signatures of probabilistic associative learning under noise and cue ambiguity and supplies trial-level measures of expectation and surprise. Exact Bayesian filtering over a small discrete state space, combined with a single local Hebbian rule, is therefore sufficient to account for these phenomena without an explicit optimization objective and without a parametric approximation of the posterior.

## Results

Through simulations, we investigated the sensitivity of our model to environmental noise, stochastic transition dynamics, and non-stationary changes in transition structure. We also tested the ability of the model to capture the learning dynamics underlying both short- and long-timescale statistical regularities in the environment. In addition, we used two experimental datasets to evaluate whether the model could reproduce behavioral effects observed across individual participants in tasks relying on probabilistic associative learning under noise and cue ambiguity respectively.

### Robustness to Noise

The first set of simulations evaluated how estimates of environmental dynamics are affected by noise (that could represent noise in the environment or noise introduced by the sensors) and the stochasticity in the environment. Figure 1 shows the directed graphs depicting transition probabilities between states from which the state sequences are drawn (left-most column, top: a deterministic sequence; bottom: a stochastic sequence). The learned dynamics at various levels of noise (an ideal sensor [i.e. no noise], low noise levels [Gaussian noise with *σ* = 0.3, centered around the state-related observation], higher noise level [Gaussian noise with *σ* = 0.5, centered around the state-related observation]) are presented in the central columns of Figure 1. It should be noted that the model is not restricted to a specific noise distribution; Gaussian noise was used in the simulations for simplicity. The last column in Figure 1 reports models trained for 45 time-points on low noise but then transitioned to a completely uninformative sensor (i.e. all observations had an equal likelihood of occurring) to test how the sensor accuracy affects the posterior beliefs. For visualization purposes we present the averaged posteriors based on the sampled states to show that the posterior beliefs inferred by the model are aligned with the sampled states except when the sequence was stochastic and the sensor turned off (bottom right panel). In that case, even though the model could learn the right transitions, its posterior probabilities diverged from the simulated ones which was expected due to the stochasticity in the state transitions (see Table 5 for the numeric values of all posteriors probabilities). Comparing internal and true dynamics also indicates that our model was able to learn the true state transitions in all cases (Figure 2). To better appreciate the effect of inference and learning, we constructed two baseline models, one model that does not update its internal representation and instead keeps its dynamics fixed at the ground truth, see 8. The second baseline model that does not perform inference and instead uses its observations to update its internal representations, see appendix Figures 9 and 10. These baseline models show that while inference on transitions that are fixed to ground truth retrieves adequate posteriors, updating internal representations without inference has a clear disadvantage in high noise regimes.

**Figure 1:**
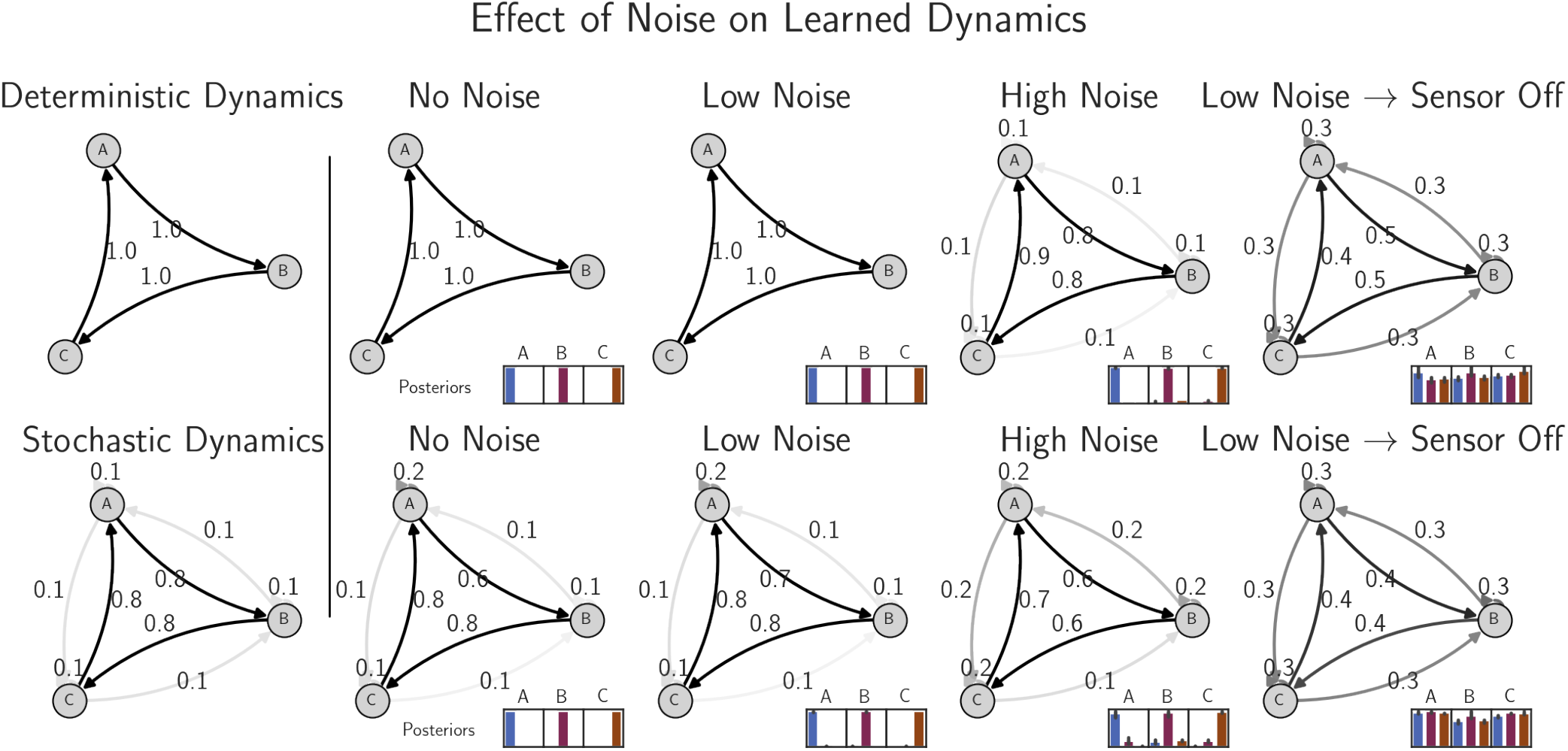
Left-most column shows the ground truth dynamics for a deterministic sequence (top) and a stochastic sequence (bottom). Numbers next to arrows denote transition probabilities. The remaining columns show in order: the learned dynamics based on an ideal sensor (i.e. no noise), sensor with low noise levels *ϵ ∼ N* (*µ*_*x*_, 0.3), higher noise levels *ϵ ∼ N* (*µ*_*x*_, 0.5), and low noise followed by an uninformative sensor. Insets next to each dynamics show the models average beliefs for each state (averaged over each occurrence of states for visualization with error bars representing 95% confidence intervals). See appendix Figures 8 and 9

**Figure 2:**
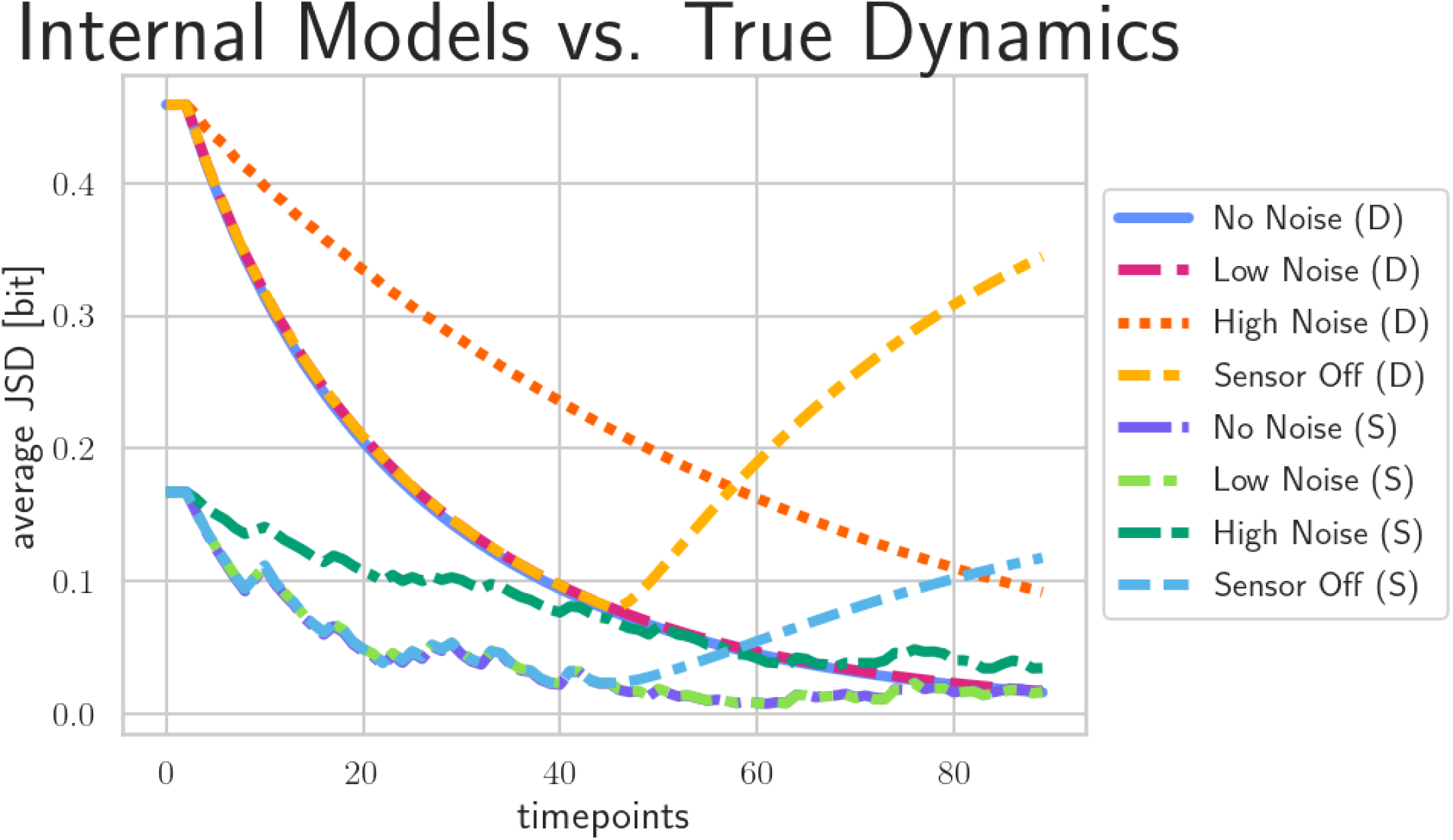
Lines summarize Jensen-Shannon divergences between the learned internal representations and true dynamics over time in simulation 1 (D: deterministic case; S: stochastic case). For a comparison with a model without inference, see Figure 10 in the appendix.

### Non stationary Dynamics

In the second simulation we tested how the model adapts to changing environmental dynamics. We sampled 200 states from an environment in which state transitions were deterministic (first half of the inputs presented to the model) and another 200 states from an environment in which state transitions were stochastic (second half of the inputs presented to the model). The model then observed the state-related information using the sensor. The results (Figure 3) show that the model quickly adapts to the true dynamics (quantified using the average Jensen-Shannon divergence between model representation and true dynamics). After the transition to stochastic dynamics, the entropy of the internal representation of the model adapts to the entropy rate of the dynamics which highlights that the uncertainty of the model mimics that of the environment. Furthermore, model predictions dynamically adapt following the transition from deterministic to stochastic dynamics, becoming increasingly uncertain and multimodal as the environment becomes more stochastic.

**Figure 3:**
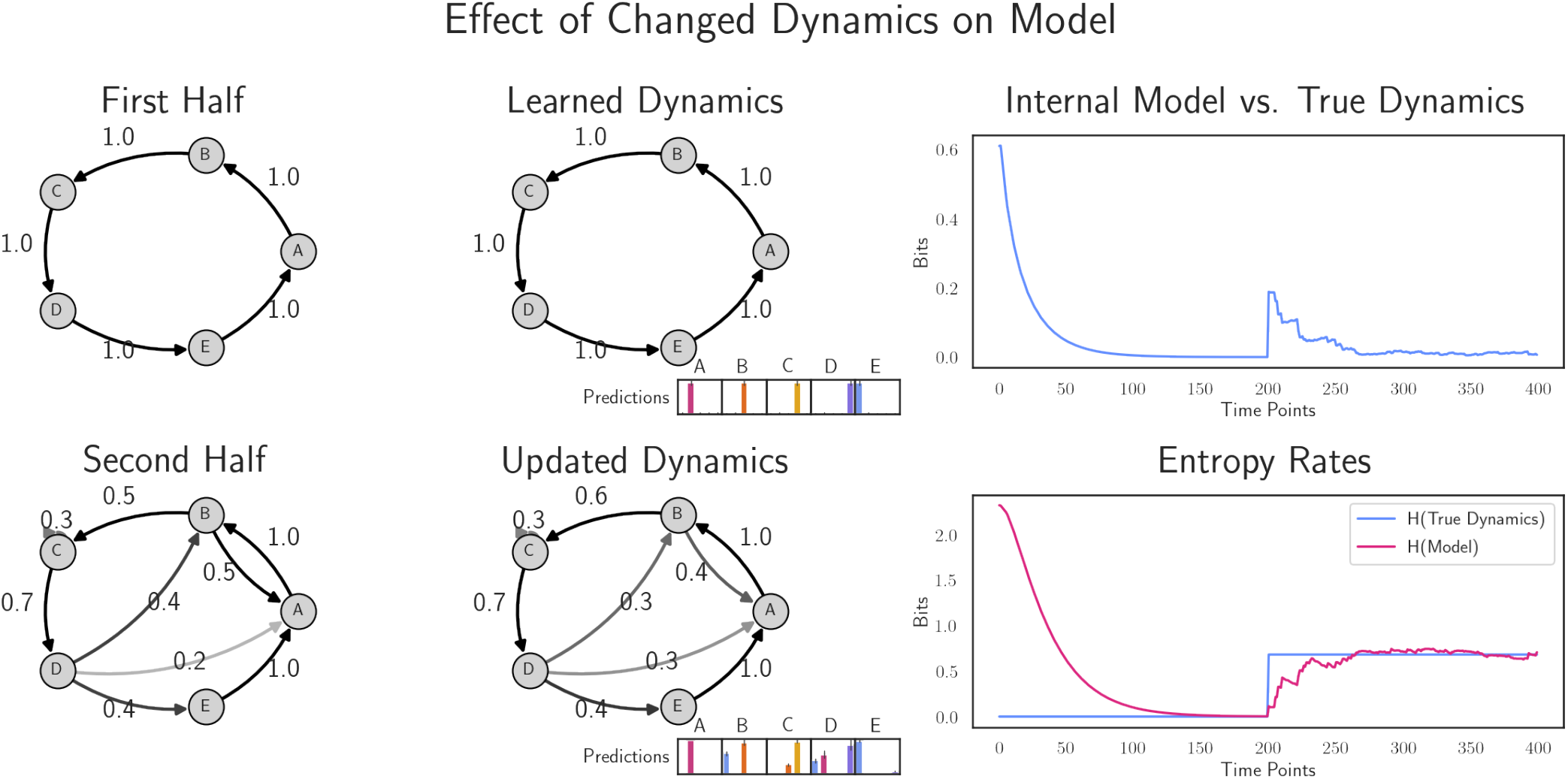
Left-most column shows the ground truth dynamics for the first half of the samples (top) and the second half (bottom). Second column shows the learned dynamics after each half. Note that the inset shows predictions rather than posteriors shown in Figure 1. In the right columns we report on the top the divergence between the internal model and the true dynamics matrix (Jensen-Shannon divergence between model representation and true dynamics) and on the bottom the entropy rates over time-points. Learning rate was set to 1.13 which was chosen based on a parameter sensitivity analysis, Figure 11.

### Learning short and long range statistical regularities

We simulated environmental transitions that follow a local-global paradigm (see Figure 4), an experimental approach that is commonly used to investigate the brain’s ability to process regularities that span multiple hierarchical levels [44, 45]. Neu-rophysiological evidence has shown that local and global statistical regularities in such paradigms are represented at different hierarchical levels in the brain, giving rise to distinct prediction error responses [45]. Computational

**Figure 4:**
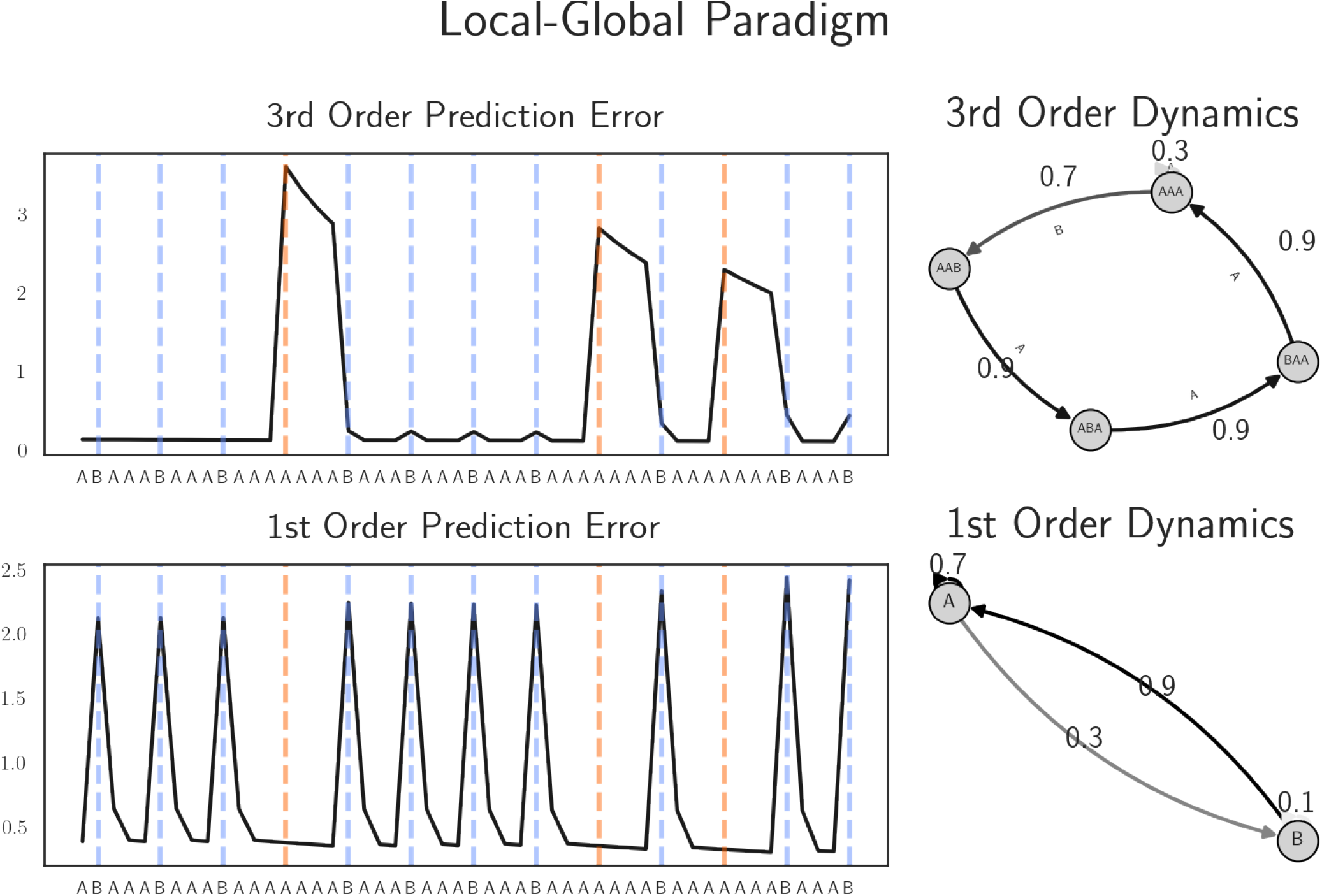
Simulated results for a hierarchical model presented with stimuli from local-global paradigm. Results on the left show errors at different timescales (top: 1 time point, bottom: 3 time points). Figures on the right show the learned transition dynamics with arrows showing the transition probabilities between posteriors.

In Figure 4 (left), we present the internal dynamics of our model for the last 50 stimuli of a sequence in which three consecutive A stimuli **AAA** are most likely followed by a B stimulus. Within this sequence, the presentation of **AAAA** represents a globally deviant stimulus in which local (short range) regularities are not violated (local standard). To capture both local and global effects we used a hierarchical version of our model and we report the error dynamics at both the lower and higher level of the model (Figure 4, left bottom and top panels respectively). At the lower level, error dynamics are affected by locally deviating (but globally predictable) presentations of B stimuli. The response to the globally predictable presentation of B stimuli is suppressed at the higher level, which instead signals the presentation of an unexpected A stimulus following **AAA**. These results provide computational evidence that local and global (expectation driven) dynamics can emerge from a hierarchical exact inference model which processes information over differing timescales.

### Modeling behavior in decision-making tasks

We used our model to capture participants’ behavioral responses in two experiments that investigated the influence of predictions on behavior (decisions and reaction times).

In Experiment 1, participants’ behavior differed depending on whether the target tone was expected or unexpected based on the preceding cue. Target tones were presented at a level relative to background noise that was calibrated during a prior staircase procedure to correspond to a detection threshold of approximately 70%, 14. Expected tones produced higher (t(35)=5.91, p¡0.05, Hedges’ g: 1.12) hit rates (mean: 77.23%, 95% CI: [73.07, 81.25]%) compared to unexpected tones (mean: 60.44%, 95% CI: [54.71 65.67]%), indicative of a cue-induced benefit in perceptual decision-making under noise. When exposed to the same stimulus sequences, SLIM reproduced the qualitative effect of cue-dependent expectations on detection performance, with higher hit rates (t(35)=58.94, p¡0.05, Hedges’ g: 14.78) for expected (mean: 83.73%, %CI: [83.38 84.07]%) vs. unexpected tones (mean: 35.26%, [33.85 36.71]%). In addition, unexpected tones elicited longer (t(35)=-6.63, p¡0.05, Hedges’ g: -1.08) reaction times (in seconds) (mean: 1.36, % CI: [1.34, 1.38]) compared to expected tones (mean: 1.22, % CI: [1.21, 1.23]), [46].

In a similar fashion, the model’s Bayesian surprise, quantified as the divergence between posterior and prior beliefs, was higher (t(35)=-88.01, p¡0.05, Hedges’ g: -19.83) for unexpected (mean: 1.66, % CI: [1.62 1.69]) than expected tones (mean: 0.42, %CI [0.4 0.45]), mirroring the direction of the participants’ reaction-time effects. We used Bayesian surprise as a proxy for the model’s reaction time, because previous behavioral works showed that Bayesian surprise predicts response times to unexpected sensory events, [47]. In the behavioral results of experiment 2, a repeated-measures ANOVA revealed a significant main effect of condition on hit rates, *F* (2, 48) = 35.18, *p <* 0.05, expected: (mean: 76.82 %, % CI: [72.21 81.29] %), unexpected: (mean:59.74 %, % CI: [54.97 64.87] %), ambiguous: (mean: 72.42 %, % CI: [68.39 76.46] %). Post-hoc pairwise tests revealed higher hit rates for expected than for unexpected (t(24)=6.08, *p*_*holm*_¡0.05, Hedges’ g: 1.34), higher hit rates for expected than for ambiguous tones (t(24)=3.62, *p*_*holm*_¡0.05, Hedges’ g: 0.39) and higher hit rates for ambiguous tones than unexpected (t(24)=6.31, *p*_*holm*_¡0.05, Hedges’ g: 1.04). Similarly, SLIM models resulted in a significant main effect of condition on hit rates, *F* (2, 48) = 1367.58, *p <* 0.05, expected: (mean: 82.24 %, % CI: [81.69 82.87] %), unexpected: (mean: 37.59 %, % CI: [35.82 39.44] %), ambiguous: (mean: 67.60 %, % CI: [66.46 68.72] %). Post-hoc pairwise tests showed higher hit rates for expected than for unexpected (t(24)=52.17, *p*_*holm*_¡0.05, Hedges’ g: 12.48), higher hit rates for expected than for ambiguous tones (t(24)=-23.98, *p*_*holm*_¡0.05, Hedges’ g: 6.04) and higher hit rates for ambiguous tones than unexpected (t(24)=27.77, *p*_*holm*_¡0.05, Hedges’ g: 7.46). Reaction times also resulted in a significant main effect, *F* (2, 48) = 39.23, *p <* 0.05, expected: (mean: 1.31 % CI: [1.27 1.36]), unexpected: (mean: 1.45 % CI: [1.39 1.5]), ambiguous: (mean: 1.38 % CI: [1.33 1.43]). Post-hoc analyses showed shorter reaction times in response to expected targets compared to unexpected targets (t(24)=-6.52, *p*_*holm*_¡0.05, Hedges’ g: -0.95) and ambiguous targets (t(24)=-5.58, *p*_*holm*_¡0.05, Hedges’ g: -0.47) and shorter reaction times to ambiguous targets compared to unexpected targets (t(24)=-6.06, *p*_*holm*_¡0.05, Hedges’ g: -0.48). Similarly, Bayesian surprises showed a significant main effect, *F* (2, 48) = 4418.69, *p <* 0.05, expected: (mean: 0.5 % CI: [0.5 0.51]), unexpected: (mean:1.67 % CI: [1.64 1.69]), ambiguous: (mean: 0.91 % CI: [0.9 0.92]). Post-hoc analyses showed shorter reaction times in response to expected targets compared to unexpected targets (t(24)=-85.87, *p*_*holm*_¡0.05, Hedges’ g: -24.91) and ambiguous targets (t(24)=-58.08, *p*_*holm*_¡0.05, Hedges’ g: -16.18) and shorter reaction times to ambiguous targets compared to unexpected targets (t(24)=-48.72, *p*_*holm*_¡0.05, Hedges’ g: -14.79). 4418.69 These results demonstrate that SLIM captures key behavioral signatures of expectation-dependent perceptual decisions and motivated further evaluation of its ability to explain decision-making behavior.

### Bayesian Modeling Analysis

To investigate whether model-derived information quantities could explain individual differences in perceptual decisions, we tested whether information quantities derived from the model were associated with participants’ perceptual sensitivity (d’) and reaction times using Bayesian hierarchical regression. All Bayesian regression models (which use different predictors described in S1.11) converged without warnings and showed satisfactory convergence diagnostics, indicating adequate sampling of the posterior distributions. We compared models containing different predictors using leave-one-out cross-validation (LOO-CV) to assess whether any information quantity provided superior predictive performance. Across all comparisons, no model showed a substantial advantage over the others. Nevertheless, we further examined the model with the highest predictive performance in each analysis. We additionally assessed model adequacy using posterior predictive checks, summarized by Bayesian p-values, where values close to 0.5 indicate that the observed data are consistent with the model’s posterior predictive distribution. In Experiment 1, among the compared predictors for sensitivity (d’), the *I*_*prior*_ model showed nominally highest predictive performance (*elpd*_*LOO*_ = − 611.257, Bayesian p-value=0.49). However, it performed comparably to *D*_*KL*_(*Posterior* ||*Prior*) (Δ*elpd* = 2.956, Bayesian p-value: 0.49), followed by *H*_*posterior*_ (Δ*elpd* = 14.456, Bayesian p-value: 0.49), *I*_*posterior*_ (Δ*elpd* = 17.334), Bayesian p-value=0.49), and *I*_*ratio*_ (Δ*elpd* = 18.323, Bayesian p-value=0.49) see Table 1.

**Table 1:** Experiment 1 LOO model comparison for d’.

|  | rank | elpd_loo | p_loo | elpd_diff | weight | se | dse | warning | scale |
| --- | --- | --- | --- | --- | --- | --- | --- | --- | --- |
| $I_{prior}$ | 0 | -611.257 | 38.345 | 0.000 | 0.696 | 26.161 | 0.000 | False | log |
| Bayesian surprise | 1 | -614.213 | 38.084 | 2.956 | 0.193 | 25.785 | 3.846 | False | log |
| $H_{posterior}$ | 2 | -625.713 | 37.606 | 14.456 | 0.000 | 24.859 | 5.869 | False | log |
| $I_{posterior}$ | 3 | -628.591 | 37.847 | 17.334 | 0.000 | 24.544 | 6.321 | False | log |
| log ratio | 4 | -629.580 | 37.676 | 18.323 | 0.111 | 24.134 | 7.315 | False | log |

We consider here *I*_*prior*_ for further analysis because it is simpler than models that use more than one probability mass (*H*_*posterior*_, *D*_*KL*_(*Posterior*|| *Prior*), *I*_*ratio*_). *I*_*prior*_ showed a positive association with d’ (*β* = 0.187, 95% HDI [0.068, 0.307]). Condition showed a negative effect on d’ (*β* = -0.365, 95% [-0.485, -0.251). The interaction between *I*_*prior*_ and condition was negative (*β* = -0.421, 95% [-0.533, -0.296]), indicating that the effect of *I*_*prior*_ differed between conditions, see Table 6. The posterior regression plot (Figure 5) indicates that the effect of *I*_*prior*_ on d’ depended on condition. Specifically, *I*_*prior*_ was associated positively with d’ for expected targets (*β* = 0.604, 95% HDI [0.408, 0.819]) and negatively with d’ for unexpected targets (*β* =−0.237, 95% HDI [-0.347, -0.111]), 7. A model comparison between condition and *I*_*prior*_ only, further showed that the *I*_*prior*_-based model showed better expected out of sample predictive performance than the condition-based model, 8.

**Figure 5:**
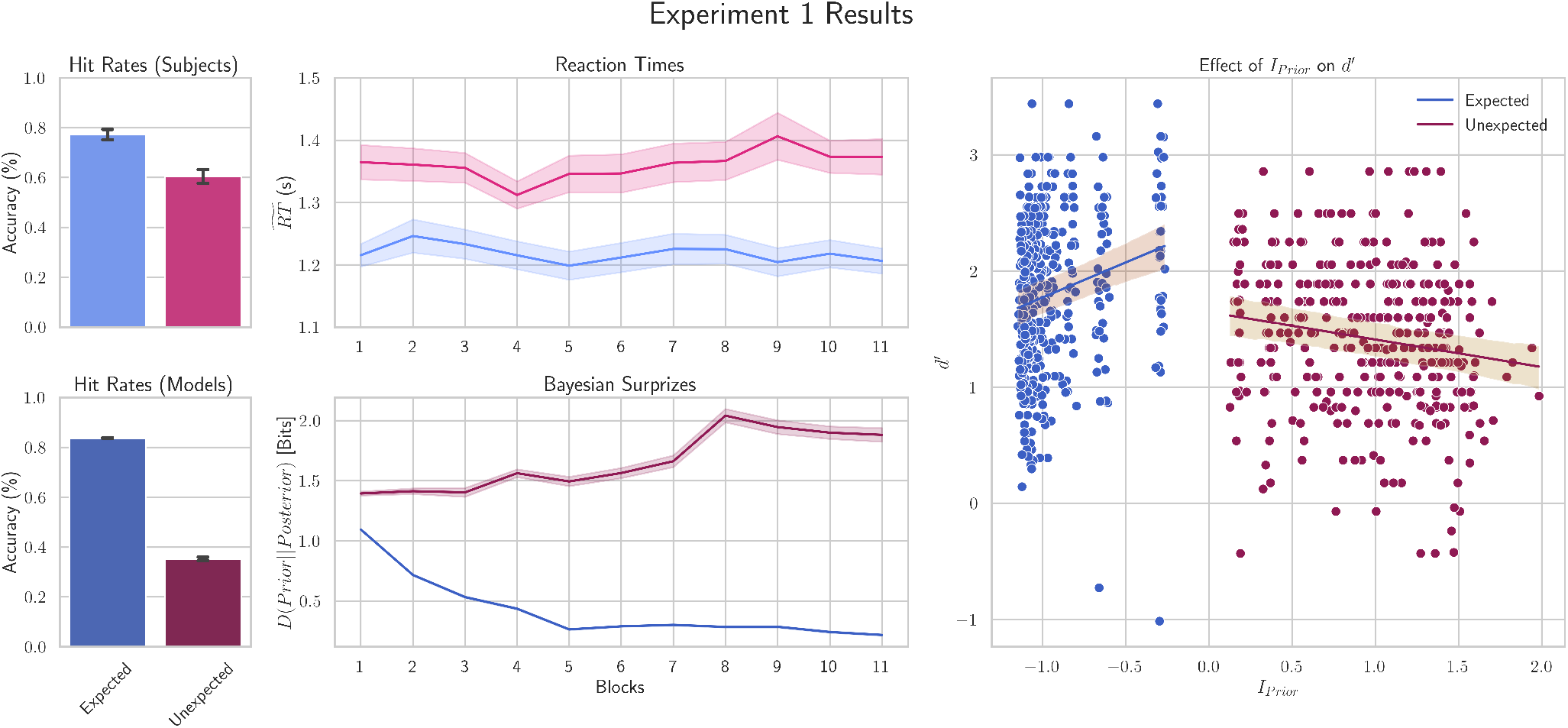
Behavioral and model results in terms of effects of expected and unexpected target tones. Left column shows hit rates for subjects on the top and for models on the bottom (blue: expected targets, red: unexpected targets). Middle column shows the reaction times to the two conditions in the top panel and Bayesian surprises in the model for the two conditions in the bottom panel. Right column shows *I*_*prior*_ plotted against d’ for individual subjects and posterior predictive regression lines with 95% highest-density intervals show that *I*_*prior*_ has a positive association with d’ for expected targets and a negative association for unexpected targets.

For reaction times, there was also no clear winning model. The highest-ranking model included (*I*_*ratio*_, Bayesian p-value: 0.5) as a predictor; however, the estimated effect was small and the posterior distribution was centered close to zero (*β* = 0.005, 95% HDI [-0.005, 0.015]). The interaction between *I*_*ratio*_ and condition was similarly small (*β* = 0.002, 95% HDI [-0.008, 0.012]), providing little evidence that *I*_*ratio*_ modulated the effect of condition on reaction times. In contrast, the condition effect itself was reliably different from zero (*β* = 0.052, 95% HDI [0.043, 0.061]), indicating that participants responded differently to expected and unexpected tones (see Tables 2 and 9). All the other models resulted in Bayesian p-values of 0.49 indicating that alternative models reproduce the data equally well.

**Table 2:** Experiment 1 LOO model comparison for RT.

|  | rank | elpd_loo | p_loo | elpd_diff | weight | se | dse | warning | scale |
| --- | --- | --- | --- | --- | --- | --- | --- | --- | --- |
| log ratio | 0 | 910.053 | 48.257 | 0.000 | 0.649 | 27.188 | 0.000 | False | log |
| Bayesian surprise | 1 | 909.179 | 48.694 | 0.873 | 0.000 | 27.335 | 2.202 | False | log |
| I_prior | 2 | 908.918 | 49.631 | 1.134 | 0.351 | 27.416 | 2.756 | False | log |
| H_posterior | 3 | 908.793 | 48.541 | 1.260 | 0.000 | 27.423 | 2.233 | False | log |
| I_posterior | 4 | 908.250 | 49.202 | 1.803 | 0.000 | 27.423 | 2.112 | False | log |

In Experiment 2, the Bayesian regression analysis resulted Bayesian p-values of 0.5 and displayed no clear winner across the different models for predicting d’, see Table 3. The highest ranking model was with Bayesian surprise (Supplementary Figure 11). For consistency with experiment 1, we picked the second highest-ranked model (*I*_*prior*_ with *elpd*_*LOO*_ = −557.384) for subsequent interpretation of regression coefficients, simple effects, and visualization of the predictive posterior regression lines. There was little evidence for predictor *I*_*prior*_ (*β* = 0.067, 95% HDI [-0.051, 0.189]) and condition (*β* = −0.065, 95% HDI [-0.162, 0.042]) on their own, but the interaction between them indicates that the outcome differed across conditions (*β* = −0.093, 95% HDI [-0.147, -0.038]), see Figure 6.

**Table 3:** Experiment 2 LOO model comparison for d’.

|  | rank | elpd_loo | p_loo | elpd_diff | weight | se | dse | warning | scale |
| --- | --- | --- | --- | --- | --- | --- | --- | --- | --- |
| Bayesian_surprise | 0 | -556.784 | 29.177 | 0.000 | 0.604 | 31.450 | 0.000 | False | log |
| x_I_prior | 1 | -557.384 | 28.991 | 0.600 | 0.392 | 31.426 | 2.413 | False | log |
| x_H_posterior | 2 | -559.924 | 28.863 | 3.140 | 0.004 | 31.149 | 2.831 | False | log |
| x_I_posterior | 3 | -560.665 | 28.472 | 3.881 | 0.000 | 30.810 | 3.184 | False | log |
| x_log_ratio | 4 | -562.355 | 28.862 | 5.570 | 0.000 | 30.871 | 3.415 | False | log |

**Figure 6:**
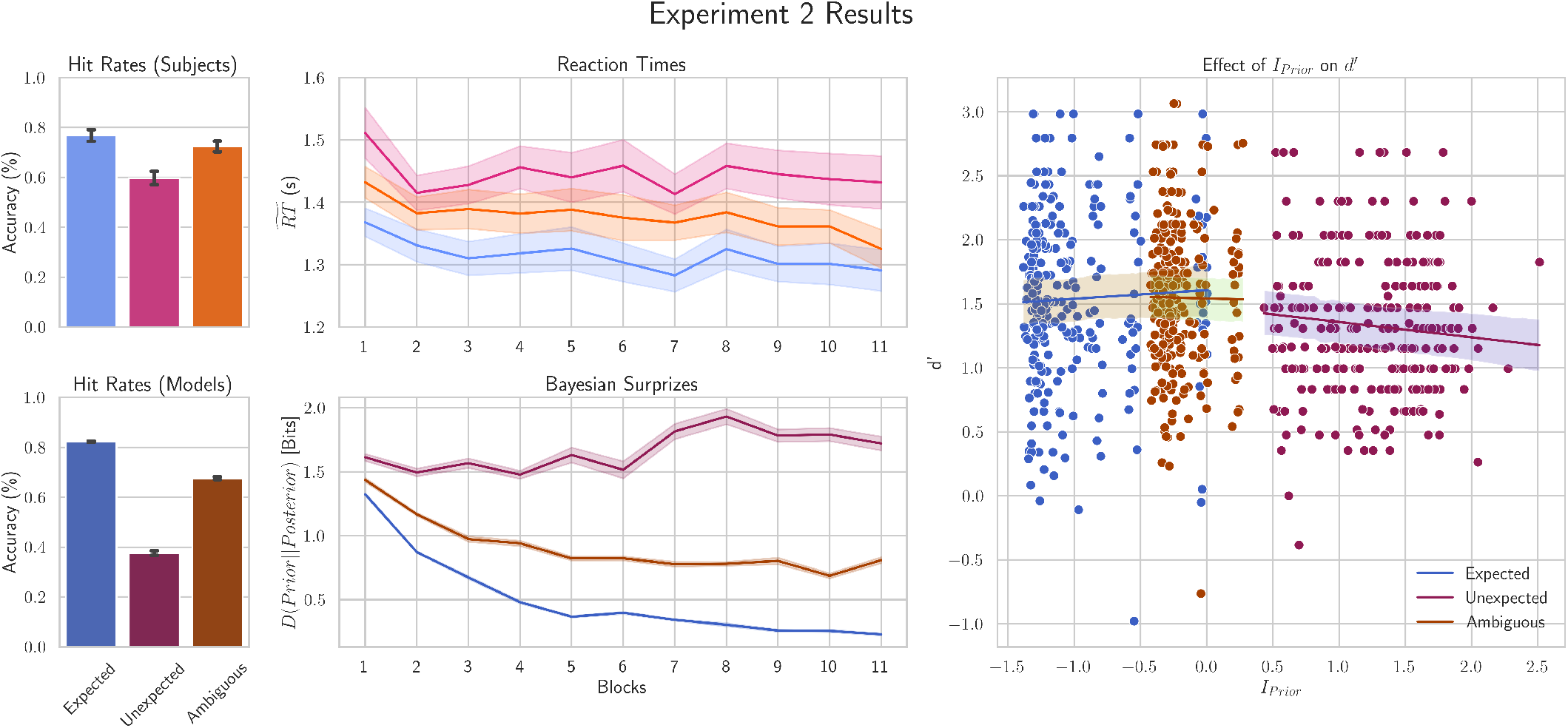
Legend as in Figure 5. Ambiguous condition is colored in orange. The right column shows predictive regression lines with 95% highest-density intervals which indicate that *I*_*prior*_ has no evidence for positive association with d’ for expected targets, no evidence for an association with ambiguous targets but a negative association for unexpected targets.

The analysis of the simple effects indicated a negative association between *I*_*prior*_ and d’ for unexpected targets (*β* = −0.120, 95% HDI [-0.203, -0.034]), weak evidence for a positive association with d’ for expected targets (*β* = 0.067, 95% HDI [-0.051, 0.189]), and no evidence for an association with ambiguous targets (*β* = −0.028, 95% HDI [-0.120, 0.055]), see Figure 6 and Supplementary Table 12.

Again, the analysis of reaction times resulted in consistent goodness of fit (Bayesian p-value=0.51) and showed no clear winner. In this experiment the highest ranking model (*D*_*KL*_(*Posterior*||*Prior*), *elpd*_*LOO*_ = 696.088) showed evidence for an effect of Bayesian surprise on reaction times (*β* = 0.031, 95% HDI [0.018, 0.042], an effect of condition (*β* = 0.024, 95% HDI [0.013, 0.034]) and interaction between Bayesian surprise and condition (*β* = −0.011, 95% HDI [-0.018, -0.004]), see Tables 4 and 13.

**Table 4:** LOO model comparison for RT.

|  | rank | elpd_loo | p_loo | elpd_diff | weight | se | dse | warning | scale |
| --- | --- | --- | --- | --- | --- | --- | --- | --- | --- |
| Bayesian_surprise | 0 | 693.385 | 30.422 | 0.000 | 0.752 | 32.884 | 0.000 | False | log |
| x_I_prior | 1 | 690.322 | 30.012 | 3.063 | 0.000 | 33.112 | 2.589 | False | log |
| x_log_ratio | 2 | 688.374 | 30.572 | 5.011 | 0.248 | 32.240 | 4.614 | False | log |
| x_I_posterior | 3 | 687.391 | 30.513 | 5.994 | 0.000 | 32.477 | 4.247 | False | log |
| x_H_posterior | 4 | 686.775 | 31.072 | 6.610 | 0.000 | 33.216 | 4.642 | False | log |

## Methods

### Model Description

SLIM assumes that the environment consists of a finite set of observable discrete states which evolve over discrete time. Crucially, the model does not have direct access to these states, but instead receives (possibly noisy and incomplete) information about these states using a sensor (see next section). As a consequence, the model learns about the environment and its dynamics and infers states based on incoming sensory information. SLIM achieves this via Bayesian filtering, to sequentially infer the most likely distribution of states and updating an internal representation that summarizes beliefs about environmental transitions. The Bayesian filter recursively infers beliefs over the current state *x*_*t*_ based on beliefs over the previous state *x*_*t−*1_ *∈ X* and the most recent observation *y*_*t*_ *∈ Y*:

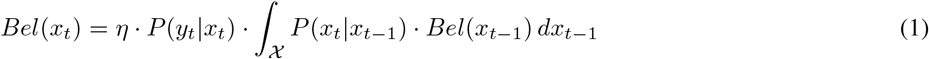

where *Bel*(*x*_*t*_) is the posterior belief at time *t* given the past trajectory of observations. In other words, Bayesian filtering calculates the probability distribution of the (unobservable) current state based on the entire sequence of previous observations,i.e. *P* (*x*_*t*_ |*y*_1,…,*t*_), see S1.1 for its derivation. *P* (*y*_*t*_ |*x*_*t*_) is a sensor model that specifies the probability of observing *y*_*t*_ *∈ Y* if the true state is *x*_*t*_ (i.e. the likelihood of an observation given a state). Note that we purposely do not characterize the domain of the observation space, because it can take different forms depending on the use case. In the simplest case *Y* = *X*, while in other cases, *Y* could be, for example, the input space to a neural network. The requirement here is that a mapping exists between *X* and *Y* and that this mapping provides information about the latent states. *P* (*x*_*t*_|*x*_*t−*1_) is the internal representation of the environmental dynamics. It describes (hidden) state transitions from *x*_*t−*1_ to *x*_*t*_. *Bel*(*x*_*t−*1_) is a recursive term that represents the belief about the previous state. *η* is a normalization term which ensures that the posterior is a proper probability distribution (i.e. that all probability masses of the posterior sum up to 1). The two terms in the integral in equation (1) can be intuitively understood from the models’ perspective as the following: *Bel*(*x*_*t−*1_) *≡* ‘*I believe this was the previous state of the environment*’ and *P* (*x*_*t*_ |*x*_*t−*1_) ‘*I* ≡ *believe the environment evolves like this*‘. The integral term thus predicts the current state distribution by marginalizing over all possible previous states (‘… *therefore this is likely the state of the environment*‘). Given our discreteness assumption, we can represent the probability distribution of a state as a vector **x** = [*x*_1_ … *x*_*d*_] ^⊺^ with *d* distinct states. The Bayesian filter then updates beliefs over the hidden states recursively as:

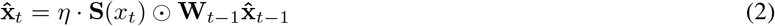

where the hat symbol denotes beliefs (posteriors), the bold font represents the full distribution rather than individual probability masses, **S**(*x*_*t*_) is the observation of the sensor model given state *x*_*t*_, and **W**_*t−*1_ is the internal representation of the hypothesized environmental dynamics prior to the observation. Note that even though we do not claim that the brain performs this exact transformation, all operations in our model including element-wise, matrix-vector, and normalization operations are biologically plausible through basic neural mechanisms [48].

### Sensor

The sensor compartment represents the model’s interface with the external world. The sensor is a model of the sensory organ (e.g. ear or eye) that receives signals from the environment (e.g. an auditory or visual stimulus) and from which a representation of the environment emerges. We frame this representation as a distribution of probabilities over the possible observations. Since our model assumes discrete states, the sensor of SLIM can be implemented as a linear mapping (a row stochastic matrix) **S** ∈ [0, 1]^*d×d*^ that provides a probability distribution of the observations given the hidden state at a specific moment in time,i.e. **S**(*x*_*t*_) = *p*(*y*_*t*_ |*x*_*t*_). Note that, for simplicity, we assume throughout this work that state and observation spaces are identical, even though our framework is not restricted to this case. Alternatively, the sensor could also be implemented as a non-linear feed-forward neural network **S**_*θ*_(*x*_*t*_), which maps a continuous stimulus (e.g. an auditory waveform) to a discrete probability distribution of observations defined over the discrete state-space of SLIM. This latter option has the benefit of ensuring that the model receives the exact same input as a subject in an empirical study and allows to study the effect that (different types of) noise in the environment have on the sensor and the learning and inference model (see e.g. behavioral experiments below). Other sensor models are also possible, in the auditory domain for example the sensory input could be represented by a cochleagram transformation [49].

### Internal Model

The internal representation **W** keeps track of environmental dynamics. We formally define this internal representation (or internal model) as a Markov transition matrix that SLIM learns based on beliefs. A Markov transition matrix **W** ∈ [0, 1]^*d×d*^ is a row-stochastic matrix defined over the finite state space *X*, but seel[50]l. Every row *i* in **W** represents the conditional probability distribution of transitioning from state *x*_i_ to state *x*_j_, i.e. 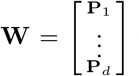. where 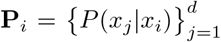 and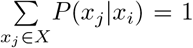. Thus, each entry in **W** represents the conditional probability mass of transitioning from state *x*_*i*_ at time *t−* 1 to state *x*_*j*_ at time *t*:

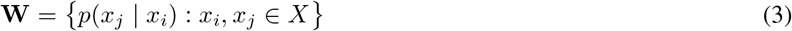

An entry in this matrix could be intuitively thought of as a hypothesis of a causal (or temporal) relationship between two states. In this formulation, we assume that state transitions are memory-less. In other words, a state *x*_*j*_ only depends on the immediate previous state *x*_*i*_, but see the section *Learning higher-order dynamics* which relaxes this assumption, enabling learning of higher-order dynamics. Note that **W** can be interpreted as a fully connected recurrent network where each cell quantifies the connection strength between state transitions.

### Learning

At each time point, the internal representation is updated using the following rule:

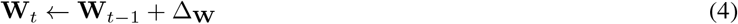

with update signal Δ_**W**_. This learning rule operates over probability distributions and is inspired by established learning mechanisms such as the ones described by [51] and [52].

To guarantee that the updated conditional probabilities (rows in the dynamics matrix) are well formed distributions (i.e. probability masses are non-negative and sum to one) and that updated transitions in the internal representation match the estimated transitions between different states, we derived a gated Hebbian learning rule with two components. The first, 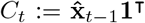, represents the context (because it considers past beliefs), and broadcasts previous state(s) onto the error term. The net effect of this matrix is always inhibitory in that it can only reduce the potential error. The second Hebbian term is 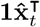 which could be thought of as a copy (broadcast) of the current state-belief, and its subtraction with the previous internal representation provides an ‘error potential’ which is inhibited by context. The update signal is therefore the context-weighted difference between the current state belief and the previous representation of the transitions **W**_*t−*1_:

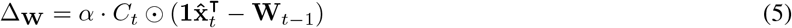

with the Hadamard operator ⊙(i.e. element-wise matrix multiplication) representing element-wise contextual weighting. This decomposition of the transition estimate has the consequence of updating the conditional distributions in **W** depending on how relevant they are within the current context *C*_*t*_. In the appendix (**Example 1**) we provide some intuition on how this Hebbian-based learning rule results in well formed probabilities and how the internal model represents transitions between states. The learning dynamics of individual cells within the internal representation of SLIM correspond to a form of exponentially weighted moving average (EWMA). In other words, within each particular transition probability estimate that SLIM learns, its update rule is equivalent to an EWMA - see S1.3. EWMAs are equivalent to discrete-time subthreshold dynamics of a leaky integrate-and-fire (LIF) neuron, [53]. Thus, learning within a SLIM model corresponds to sub-threshold membrane dynamics of a network of discretized LIF units which integrate input (posterior probability masses) and leak previous memory (probability masses of previous hypotheses). While standard time-varying EWMAs are guaranteed to asymptotically converge to a constant input if *α*_*t*_ ∈ (0, 1] and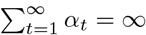, the error in the estimate of a stationary dynamics matrix can be shown to be bounded using the SLIM update rule (see S1.6. The learning rate governs how fast the model is updated at each time step. In SLIM, the learning rate can be assumed as fixed or optimized in different ways. SLIM is Lipschitz continuous with respect to its learning rate when it is restricted in *α* ∈ [0, 1], S1.5. This enables global Lipschitzian optimization of the learning rate (and potentially other variables like hierarchical weights and sensor models) which provides an alternative to gradient-based optimization.

### The Step Function

At each time-step, the sensor **S** observes new information (a snapshot) related to the current state and transforms it into a probability distribution of possible states capturing the uncertainty about sensory inputs. Concurrently, the model predicts the current state based on the previous internal representation of state transitions and its belief about the previous state. This prediction is probabilistic, and thus captures the internal models’ uncertainty about the environment. Finally, the model is updated using the update signal (the difference between the current state [matrix] and the previous representation of dynamics) weighted by the current context. The context weighs the transition probability distributions (rows of the internal representation) based on their recency, such that only relevant distributions are updated.

The following pseudo-code summarizes how SLIM operates in the non-hierarchical case:

#### Algorithm 1

Simultaneous Learning and Inference

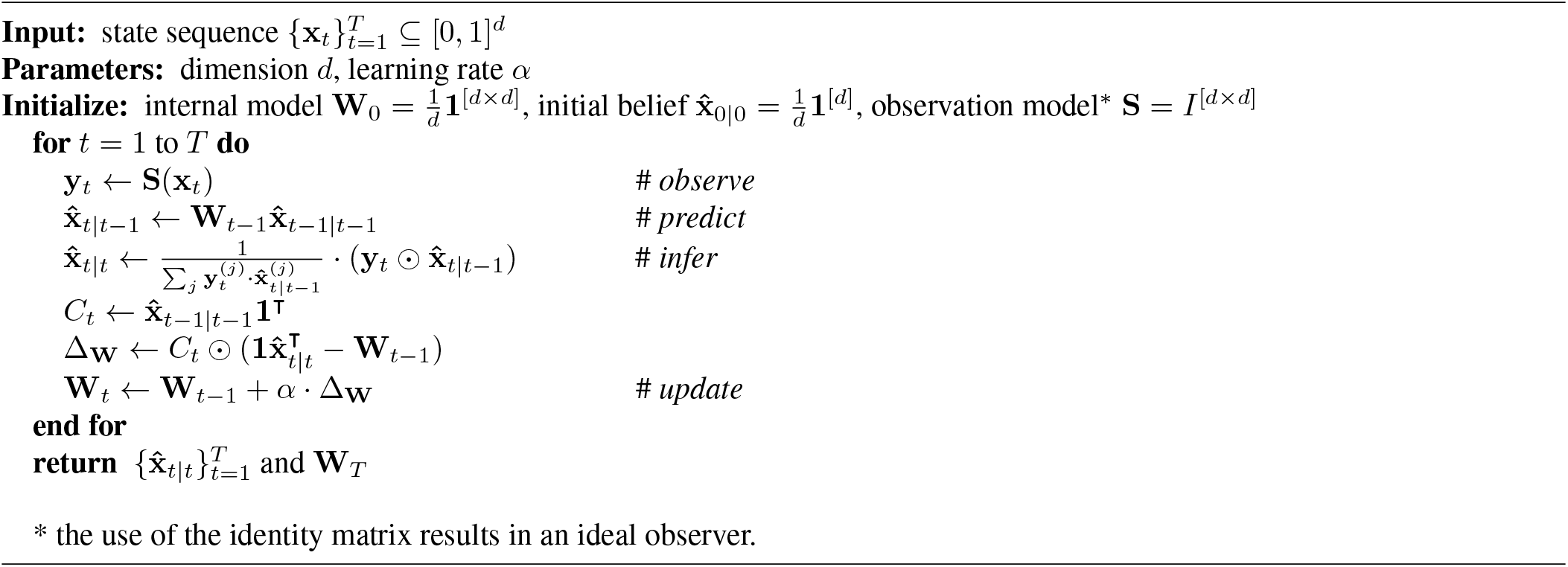

### Learning Higher-Order Dynamics

The human cortex is organized within a hierarchy of temporal receptive windows, [42]. In a SLIM model, the sensory and inference compartments form a module that can be stacked to build a hierarchal model capable of learning environmental dynamics that span longer time-scales. The Bayesian filter hinges on the Markov assumption to produce optimal belief updates. To fully characterize the past, we combine multiple previous beliefs (rather than the one immediate previous belief) to form a Higher-order Markov matrix. For this purpose, we parametrize our learning rule using a free parameter that represents the time-scale *k*:

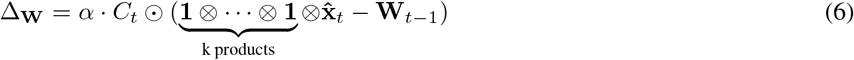

with 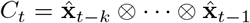. In the case of *k* = 1 the update rule reduces to the original first order update rule up to the context where in the first order rule, the previous state belief is expanded into a matrix using the **1** vector to broadcast the belief onto the previous internal representation.

### In Silico Experiments

We performed a set of simulations to test the robustness of SLIM under environmental noise, stochastic transitions, and transitions within a non-stationary environment. We also test the ability of the model to learn both short and long term statistical regularities in the environment.

### Stationary World Dynamics

To test whether SLIM can simultaneously learn meaningful representations and perform accurate state inference under different levels of sensory and environmental uncertainty, we first evaluated the model in controlled environments with known transition dynamics. The first set of simulations considered deterministic transitions between 3 states *{A, B, C}* with (*A → B → C → A→* …) at 4 different levels of constant noise in the input (no noise, low Gaussian noise, high Gaussian noise, and low Gaussian noise followed by maximum noise). The Gaussian noise was added to the discrete cells of the sensor matrix (row-wise probability distributions over possible states) by applying a Gaussian filter to an identity matrix with specified standard deviations and normalizing to get a proper probability distribution in each row. We also considered the case of stochastic dynamics (between the three states) and consider the same 4 noise levels. In both cases (deterministic and stochastic transitions) the environment is considered as stationary (i.e. not changing over time). We trained eight model instances (one for each dynamics type and noise level) that learn a 1st order transition matrix (i.e. no hierarchy). We analyzed the learned representations and posterior distributions to test whether SLIM models can learn meaningful representations and whether they produce accurate beliefs. All models were trained for a duration of 90 time points (30 repetitions per state) with the learning rate set to 0.1.

### Nonstationary World Dynamics

To test whether SLIM can adapt its internal model to changing environmental dynamics, we evaluated its ability to track non-stationary transition structures in a second set of simulations. We considered the transition from a deterministic environment with transitions between 5 states to a stochastic one. To quantify our model’s ability to flexibly adapt to changes in the environment, we used the Jensen-Shannon divergence (JSD). The JSD is a symmetric and normalized measure of divergence between two distributions. It is related to the better known Kullback-Leibler divergence and is defined as 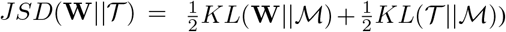, with true transition probabilities of the environment *T* and mixture distribution 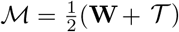. In practice, we calculated the average JSD over the different states of the environment. We also analyzed whether the learned internal representation can capture the information entropy of the environment to show whether the entropy of the internal model mirrors that of the environment.

### Local-Global Paradigm

To test our model’s ability to capture information over longer time scales, we built a hierarchical SLIM model and presented it with a stimulus sequence with both short and long term statistical regularities. We used stimuli generated using a canonical local-global paradigm [44]. This paradigm contains two types of regularities, a local (short-time) regularity (e.g. after an **A** stimulus there is a high likelihood of hearing another **A** stimulus) and a global (long-time) regularity (e.g. after **AAA** stimuli, there is a high probability that the next stimulus is **B**). This paradigm is typically used to investigate hierarchical prediction error processing [45, 54]. The violation of the local regularity (i.e. presenting a stimulus **B** after three **A** stimuli) elicits a local prediction error while violating the global regularity (i.e. presenting stimuli **AAAA** whereas the most probable stimuli are **AAAB**) is used to elicit prediction errors in response to violations of predictions that integrate information over longer time frames. To enable our model to learn local and global regularities, we built a two-level (hierarchical) model. The lower level was set to learn first-order dynamics, and the higher level was set to learn third-order dynamics (i.e. accounting for 3 time-points in the past). Instead of direct sensor-driven observations, the higher level received as input the posterior belief of the lower level. The lower level received predictions from the higher level, which were combined with the predictions from the lower level using a weighted average (.7 *×local −prediction* + .3 *× global− prediction*). Both levels were set at a learning rate of 0.2. Note that while in these simulations the weights for the combination of lower and higher level predictions as well as learning rates were fixed, they are hyperparameters that can be optimized (and cross-validated) when modeling empirical results. We present one way of optimizing such hyperparameters in the appendix and leave applications to future work. We trained the model over 200 time points and consider the case in which the sequence is formed by **AAAB** sub-sequences (i.e. a locally deviating sequence is the global standard) and **AAAA** served a test of sensitivity to deviation of the global regularity.

### Behavioral Experiments

We tested the ability of our model to reproduce empirical results in two behavioral auditory experiments. The first experiment investigated the inferential nature of perception and decision-making under noise [46]. The second experiment was built to extend the first in order to additionally investigate the effect an ambiguous cue (i.e. context) can have on processing trailing targets.

### Ethics Statement

Participants in experiment 1 were recruited between 03.09.2024 and 10.06.2025. All participants provided written informed consent before participation. The study was approved by the local ethics committee (CMO Arnhem-Nijmegen, Radboud University Medical Center, “Imaging Human Cognition”, CMO 2014/288, CCMO protocol NL45835.091.13, accessible at https://onderzoekmetmensen.nl/en/trial/55541. The study followed the principles expressed in the Declaration of Helsinki. Participants in experiment 2 were recruited between 01.05.2025 and 28.05.2025. All participants provided written informed consent before participation. The study was approved by the Ethics Review Committee for Psychology and Neuroscience (ERCPN approval code OZL-232-01-01-2021) at Maastricht University, following the principles expressed in the Declaration of Helsinki.

### Experiment 1: Probabilistic Associative Learning under Noise

In experiment 1, 40 participants with normal hearing and no history of neurological or psychiatric disorders were tested on a probabilistic auditory associative learning task. The experiment was conducted at the Donders Institute for Brain, Cognition and Behavior (Nijmegen, The Netherlands) and was approved by its ethical committee. In what follows, we provide relevant experimental details for the analyses we conducted here. All remaining information can be found in the original preprint [46]. The stimuli consisted of four pure sinusoidal tones with pitches 300, 848.53, 2400 and 6788.23 Hz (1.5 octaves apart), sampled at 48000 Hz. The tones had a duration of 50 ms, including 5 ms rise and 5 ms fall ramps. To remove inter-subject differences in terms of tone-detection thresholds, each participant completed two adaptive staircase sessions before the main task, one for each target tone (300 and 6788.23 Hz). On each trial, the target was presented in continuous white noise, and participants had to detect and identify it. Stimulus intensity was adjusted with a 1-down/1-up rule and unequal steps (down-step = 0.80 × up-step), i.e. a weighted up-down staircase, [55, 56]. This procedure tracks a point slightly above 50% correct on the staircase task. The resulting intensity for each target was then used in the main task. The step size ratio of 0.80 was chosen through piloting, so that the hit rates in the subsequent main task were around 70% at these intensities.

The main task consisted of 11 blocks, each consisting of 110 trials. During each trial, participants were presented with a pair of tones, a cue tone (848.53 or 2400 Hz) followed by a target tone (300 or 6788.23 Hz). All tones were presented together with a constant (white) noise and with a 350 ms gap between cue and target. The inter trial interval between two consecutive trials was between 1750 and 4250 ms. Before the first block, one cue-to-target pairing was selected randomly, and was kept constant throughout the experiment. The first 10 trials of each block had increased intensity for the target tones (compared to the intensity that was determined using the staircase procedure) to ensure that participants heard the cue-target relationship. These ‘easy’ trials were removed from the analysis. The 100 subsequent trials used the intensity for the target tone that was determined during the staircase. The cue tones were always presented at a clearly audible level. In 75% of the trials each cue was followed by one of the two targets which we refer to as expected target. In the remaining trials the cue was either followed by the other target (18%, unexpected target) or an omission (7%, omitted target). The task of the participants was to respond to the target tone as fast as possible by indicating what they heard (target low, target high, no sound). Participants’ data was excluded from the analysis if they reached a sensitivity index *d*^*′*^ *<* 1 for both the expected and unexpected conditions. As a result we analyzed data from 36 participants (24 female, 12 male, average age 22.41).

### Experiment 2: Learning under Cue Ambiguity

Experiment 2 was designed to test the effect of an ambiguous cue on the processing of targets. We used a similar procedure to Experiment 1 (including staircase sessions, trial construction and inter trial time in the main experiment), but we adapted the design by adding a third cue (1427.05 Hz) that was ambiguous or uninformative with respect to the target that follows it - i.e., each target was presented with equal probability following this additional cue. We recruited 30 healthy participants (25 female, 5 male, average age 22.06). None of the participants of Experiment 2 also took part in Experiment 1. All data for Experiment 2 were collected at the faculty of psychology and neuroscience of Maastricht university and approved by its ethical committee. Participants were instructed identically as to Experiment 1. After using the same criterion as in Experiment 1 (excluded from the analysis if they reached a sensitivity index *d*^*′*^ *<* 1), we analyzed data from 25 participants (20 female, 5 male, average age 22.04). The experimental code was also adapted from Experiment 1 and was written entirely in Matlab (Psychtoolbox, [57, 58]). The experiment lasted approximately 1.45 hours and contained 11 blocks with 127 trials per block. Each block began with 12 practice trials (4 per cue) where the trailing target tone was presented at 10 dB above the intensity determined during the staircase session, resulting in 115 trials for analysis. Similar to the previous experiment, in each block, 75% of the targets that followed the 848.53 and 2400 Hz cues were expected, 17% were unexpected, and at 8% were omitted. The ambiguous cue (1427.05 Hz) was followed by one of the two targets with a 46% equal probability. In the remaining 8% of the trials it was followed by an omission. This resulted in 58 expected targets, 34 unexpected targets, 14 ambiguous targets, and 9 omissions in each block.

### Model Instantiation

We fitted separate SLIM models for each participant using the trial-by-trial stimulus sequences presented in each experiment. We designed a sensor model (a multiscale convolutional neural network) that receives as input the sound waveforms (i.e. the combined tone [cue or target] and noise) that were presented to the participants and we trained the model to map the noisy waveforms into a probability distribution over possible states. We built this network by concatenating 1d convolutional layers of kernel sizes (161, 57, 33, 21, 7) to roughly match the periods of the different frequencies that were used in both experiments. This was followed by rectified linear units and maximum pool to focus only on the most salient features. The final layer was a fully connected layer to transform the representation into the probability of hearing one of the possible tones (i.e. 5 for experiment 1 and 6 for experiment 2). The training data used the 6 waveforms which were identical to the waveforms used during the behavioral experiments. The network was trained with the objective of minimizing the relative entropy (KL divergence) between the output logits and the true labels. During model deployment, we applied a softmax function in order for the output logits to mimic a proper probability distribution. Since no more than 6 samples were used, we highly regularized the network to learn robust representations. We did this by augmenting the samples during training (by adding uniform noise) and by adding a weight decay penalty on the gradients.

Participants in the experiment undergo a staircase procedure to determine the level of the target tone that results in a 70% detection threshold. To mimic outcome of the staircase procedure in the sensor, we varied the sound intensity (relative to the background white noise) of the target tones and calculated the classification accuracies for each target separately at each intensity level (50 times). When presenting the tone pairs (cue, target) to SLIM, we selected the level for each target tone that resulted in approximately 70% correct classification accuracy, see 13.

### Analysis

SLIM’s performance in predicting behavioral outcomes (*d*^*′*^ and *reaction times* was assessed using Bayesian hierarchical regression implemented using PyMC [59]. We evaluated whether different model predictors (information theoretic quantities derived from SLIM) better explained the behavioral performance by fitting separate Bayesian models for each predictor; see supplement S1.11 for an explanation of the different predictors. We z-scored all information theoretic quantities to place them on a common scale and facilitate the comparison of the regression coefficients. We modeled the outcomes as 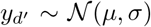 and *y*_log *RT*_ *∼ t*_*v*_ (*µ, σ*) where *µ* = *a*_0_ + *a*_*s*[*i*]_ + *β*_condition_ *c*_*i*_ + *β*_*x*_ *x*_*i*_ + *β*_interaction_ *c*_*i*_*x*_*i*_. Each model thus included the categorical condition value *c* and the predictor of interest *x* and we estimated a global intercept *a*_0_, subject specific random intercepts *a*_*s*[*i*]_, a fixed effect (slopes) of condition *β*_condition_, and the effect related to the predictor of interest *β*_*x*_, as well as the interaction between the predictor of interest and the condition *β*_interaction_. In all models, we used identical standard normal priors for the global intercept and regression coefficients while the standard deviation of the participant-specific intercept was modeled with a half (non-negative) normal. Models were estimated using the built in No-U-Turn Sampler (NUTS) of PyMC. We used four Markov chains with 1000 tuning iterations followed by 1000 posterior samples per chain. We set the target acceptance probability to 0.95 to improve sampler stability. We assessed convergence using the rank-normalized statistic 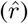 and effective sample sizes (ESS). Models were compared using approximate leave-one-out cross-validation (LOO-CV), with expected log predictive density (elpd) used to assess predictive performance. Absolute model calibration was evaluated using posterior predictive checks (PPC). Regression coefficients are reported as posterior means together with 95% highest-density intervals (HDIs). For models exhibiting significant interactions, conditional (simple) slopes were computed for each experimental condition from the posterior distribution.

## Discussion

We propose a formal predictive model that captures computational principles underlying our brains’ capability to adaptively infer environmental states (and their probabilistic transitions) from noisy inputs. By combining Bayesian filtering with Hebbian learning of belief dynamics we provide a framework for exact inference using unsupervised local learning mechanisms. This unified framework allows simultaneous learning and inference by optimally combining the input (sensory information) with predictions that result from pushing forward prior beliefs through a dynamically updated internal representation of environmental transitions.

Using both simulations and real behavioral data, we show that exact Bayesian inference and online learning of a Markov process is an appropriate approach for modeling perception in noisy and uncertain environments within the Bayesian brain hypothesis. First, we performed a set of in silico simulations to validate that SLIM is robust to environmental noise and can successfully rely on its internal predictions even when sensory information becomes uninformative (see Figure 1 top right). This ability parallels established approaches such as Partially Observable Markov Decision Process (POMDP), where robust inference is achieved by maintaining beliefs over hidden states and integrating sensory evidence with predictions about environmental dynamics [12].

In real life scenarios, the environment may undergo changes in event probabilities reflecting changes in contextual information. For example, an animal may learn that a particular location is reliably associated with food, but later experience a change in the environment that makes this association uncertain. In such situations, previously reliable predictions must be updated to reflect increased uncertainty in the underlying dynamics. Prior work has shown the hippocampus to be sensitive to the statistical uncertainty (i.e. the entropy) of the environment, [60]. We investigated this ability by exposing SLIM to a transition from deterministic dynamics, where each state was reliably followed by a single successor state, to stochastic dynamics in which multiple successor states became possible. These simulations demonstrated the flexibility of SLIM’s non-parametric representation. Predictions that were initially concentrated around a single likely state became distributed across multiple possible states as the environmental dynamics became uncertain, see insets in Figure 3. This ability to dynamically adjust the shape of the inferred state distribution distinguishes SLIM from approaches that rely on fixed parametric assumptions about the posterior distribution, such as unimodal Gaussian approximations [5, 6, 13, 61, 62]. Under such assumptions, uncertainty is typically represented by changes in variance around a single mode, limiting the ability to represent situations in which multiple interpretations of sensory input are simultaneously plausible. For example, an ambiguous stimulus with multiple valid interpretations may be represented by a single averaged estimate rather than a distribution capturing multiple possible states. While alternative parametric families could address this limitation, selecting a specific distributional form introduces an inductive bias about how uncertainty should be represented by the brain. Sampling-based approaches avoid this constraint by representing posterior distributions through samples rather than assuming a fixed analytical form [10, 27, 30, 63]. For example [27] showed that recurrent spiking neural networks can implement Bayesian inference by performing Markov chain Monte Carlo (MCMC) sampling. The stochastic spiking dynamics, together with biologically realistic properties such as refractory periods and recurrent interactions, allow the network to sample from a target probability distribution over network states, thereby representing posterior uncertainty over time. [10] extended this sampling framework to enable dynamic inference (Bayesian filtering) of changing environments via biologically plausible mechanisms. SLIM complements these approaches by providing a discrete, non-parametric framework that enables exact Bayesian inference without requiring sampling approximations.

Beyond its robustness to noise and its ability to flexibly adapt to changing contextual dynamics, we investigated whether SLIM can learn not just local statistical regularities, but also ones that span longer timescales. Environmental regularities are often structured across different temporal scales: short-term dependencies may capture relationships between individual events, whereas longer-term dependencies may capture higher-order sequential patterns. These statistical regularities are common in many naturalistic scenarios including speech and music [39, 64, 65]. Neurophysiological evidence suggests that the brain is capable of extracting and representing statistical regularities across multiple temporal scales (see e.g. [40, 41, 43, 44]). While classical mismatch negativity studies have provided strong evidence for predictive processing in the primary auditory cortex, neural responses to local and global expectation violations in sequences of auditory regularities have been shown to be reflected in broader cortical networks including the prefrontal cortex, [44]. These patterns can be attributed to the differences in temporal integration timescales, where late regions in the cortical hierarchy operate on longer timescales compared to hierarchically earlier regions. Consistent with this perspective, we showed that a hierarchical SLIM model consisting of two levels that learn transition statistics over different timescales can reproduce local and global prediction error signatures. These effects are in line with previous computational work showing local mismatch negativity effects emerging from biologically plausible spiking neurons, [66], and local/global effects in quantitative modeling, [67]. The hierarchical organization of SLIM resulted in stable internal representations while allowing the model to adapt to regularities occurring at different temporal scales, see figure 4 and supplementary figure 12.

Aligned with the view that perception is shaped by expectations, [46] examined the inferential nature of perception by testing how expectations influence conscious auditory perception during a challenging decision-making task under constant background noise. A second experiment extended this paradigm by additionally testing how ambiguous contextual cues influence the processing of subsequent target stimuli. To evaluate whether SLIM captures these expectation-dependent effects, we presented the model with the same stimulus sequences experienced by participants and compared model decisions, defined as the maximum of the posterior estimate of the inferred state, with human behavioral responses. Both participants and models demonstrated remarkably similar performance across conditions, achieving the highest hit rates for expected targets, moderate rates for ambiguous targets (close to the detection accuracy), and the lowest for unexpected targets. These results support both the Bayesian brain hypothesis and predictive processing frameworks, indicating that prior expectations dynamically bias perceptual decision making. These results are consistent with a body of computational work showing that expectations influence perceptual decisions by shaping the interpretation of noisy sensory evidence. To better appreciate the uncertainty in the observation that is passed from the neural network, we report a summary of the vectors of probability masses and logits for each state, see supplementary figure 14. Specifically the probability distributions for *T*_*low*_ and *T*_*high*_ in top-right heatmap highlight the uncertainty in the observation that is passed by the sensor. Bayesian observer models have demonstrated that combining prior beliefs with sensory likelihoods can account for systematic biases in perception under uncertainty [2]. Similarly, predictive coding and active inference frameworks propose that perception emerges from the interaction between sensory evidence and predictions generated by a hierarchical generative model [4, 6]. However, many implementations of these frameworks assume that the relevant generative model or prior structure is already available and focus primarily on inference within this model. In contrast, in SLIM expectation-dependent perceptual biases emerge from a system that simultaneously learns environmental transition statistics and performs Bayesian inference. Thus, SLIM provides a mechanistic account of how expectations underlying perceptual decisions can be acquired and updated through experience without requiring externally specified priors. Having fitted SLIM to data from each experiment, we derived *I*_*prior*_, i.e. the surprisal of the presented target under the model’s prior beliefs. Expected targets produced lower surprisal than unexpected targets, with ambiguous targets falling between these two conditions. This follows directly from the experimental structure, as the prior probability of an expected target is higher than that of an unexpected target. Our Bayesian regression analyses revealed an interaction between condition and *I*_*prior*_. For unexpected targets, lower surprisal (i.e., targets that became more predictable under the learned environmental statistics) was associated with higher discriminability. Conversely, for expected targets, higher surprisal was associated with higher discriminability in Experiment 1. Thus, under noisy conditions, unexpected events that are less surprising according to the learned model are easier to distinguish from background noise, while expected events that deviate more from their predicted probability are also detected more accurately. Although this pattern may appear contradictory, the absolute level of surprise differs substantially between conditions, even highly surprising expected targets remain less surprising than typical unexpected targets. One possible explanation for the reduced discriminability of highly predictable expected targets is reduced attention or overexposure effects. Consistent with this possibility, participants showed a decrease in (d’) across experimental blocks in Experiment 1 [46]. However, the current results do not provide sufficient evidence to distinguish between these potential mechanisms. Interestingly, the information measure used here captures only the surprisal of the observed event under the prior distribution rather than the full divergence between prior and posterior distributions typically considered in formulations of Bayesian surprise. This suggests that the information content of the predicted event itself may be sufficient to explain variations in perceptual sensitivity, without requiring a comparison of the complete prior and posterior distributions.These findings can be interpreted in the context of existing computational models of predictive processing. Prediction error is commonly formalized as a mismatch between sensory observations and prior expectations. Responses to these mismatches have been investigated using different computational measures of surprise, [4].

Shannon surprise quantifies the unexpectedness of a single event under a predictive distribution, whereas Bayesian surprise quantifies the change in distribution induced by that event and is computed as the KL divergence between posterior and prior beliefs. Both quantities have been proposed as candidate signals underlying perceptual inference and learning, [68–72]. Our results suggest that, at least for the present task, a simpler information-theoretic quantity—the Shannon surprisal of the observed event under the model’s prior beliefs—captures a substantial proportion of the variance in perceptual sensitivity. This raises the possibility that behavioral performance in noisy perceptual decisions may depend primarily on the predictability of the observed event itself, rather than on the complete belief update induced by that event. Whether this extends to more complex inference problems remains an important question for future work.

### Limitations and Future Directions

Several limitations of the current implementation provide promising directions for future work. First, SLIM assumes a pre-defined and fixed set of discrete latent states. In natural environments, however, the relevant latent states are rarely known a priori and must instead be discovered through experience. Addressing this challenge would require extending both the sensory representation and the internal state representation to allow the introduction of novel states. Although this lies beyond the scope of the present work, recent developments based on infinite semi-Markov models provide one possible approach for extending transition models in this direction [73]. Second, in the hierarchical implementation of SLIM, higher levels learn statistical regularities that partially overlap with those already acquired at lower levels. While this allows each level to independently construct representations over different temporal scales, it is unlikely to be computationally or metabolically optimal. Future work should investigate more efficient mechanisms for sharing learned structure across hierarchical levels while preserving the advantages of distributed inference. A further limitation is that SLIM relies on explicit state representations. Although this enables exact Bayesian inference, it limits the model’s ability to generalize previously learned transition structures to novel but structurally similar environments. Likewise, when environmental dynamics change substantially, previously learned transition statistics may be overwritten rather than reused. More flexible state representations could therefore improve both transfer learning and continual learning while maintaining the probabilistic inference framework developed here. Finally, the present work focused on environments with relatively small state spaces, where exact Bayesian filtering remains tractable. In many real-world settings, however, the number of latent states may become prohibitively large (or infinite in continuous spaces). In such cases, approximate Bayesian inference methods, including particle filtering, provide a natural extension by representing posterior beliefs through samples rather than explicit probability distributions. Despite these limitations, SLIM demonstrates that exact Bayesian inference and biologically plausible local learning can be combined within a unified framework for perception under uncertainty while requiring relatively few inductive assumptions. Future work will focus on fitting the model to electrophysiological and neuroimaging data, with the long-term goal of explaining individual differences in healthy cognition and neuropsychiatric disorders. To facilitate further research, we also aim to release an open-source Python implementation of SLIM, enabling researchers to test hypotheses related to predictive processing, exact Bayesian inference, and unsupervised learning without assuming a predefined family of probability distributions.

## Acknowledgments

We would like to thank Giancarlo Valente for his advice on statistical tests. The authors thank all members of the lab for their support.

## Author contributions

Conceptualization, F.D.M., M.E., and M.S.; methodology, M.E., M.S., F.D.M., and Y.V.; software, M.E.; investigation, M.E., Y.V., and P.J.; Formal analysis, M.E.; writing-–original draft, M.E.; writing-–review & editing, F.D.M, M.S., Y.V., and R.A.; funding, F.D.M.; supervision, F.D.M. and M.S.

## Materials & correspondence

Correspondence and material requests should be addressed to Mahdi Enan.

## Data availability

All data will be made available on publication. Private access to the data will be provided to editors and reviewers during peer review.

## Code availability

The model and analysis code used in this work are available at: https://github.com/mesoScopic-Computational-AuditioN-lab/.

